# Assessment of an organ-specific *de novo* transcriptome of the nematode trap-crop, *Solanum sisymbriifolium*

**DOI:** 10.1101/256065

**Authors:** Alexander Q Wixom, N Carol Casavant, Joseph C Kuhl, Fangming Xiao, Louise-Marie Dandurand, Allan B Caplan

## Abstract

*Solanum sisymbriifolium*, also known as “Litchi Tomato” or “Sticky Nightshade,” is an undomesticated and poorly researched plant related to potato and tomato. Unlike the latter species, *S. sisymbriifolium* induces eggs of the cyst nematode, *Globodera pallida*, to hatch and migrate into its roots, but then arrests further nematode maturation. In order to provide researchers with a partial blueprint of its genetic make-up so that the mechanism of this response might be identified, we used single molecule real time (SMRT) sequencing to compile a high quality *de novo* transcriptome of 41,189 unigenes drawn from individually sequenced bud, root, stem, and leaf RNA populations. Functional annotation and BUSCO analysis showed that this transcriptome was surprisingly complete, even though it represented genes expressed at a single time point. By sequencing the 4 organ libraries separately, we found we could get a reliable snapshot of transcript distributions in each organ. A divergent site analysis of the merged transcriptome indicated that this species might have undergone a recent genome duplication and re-diploidization. Further analysis indicated that the plant then retained a disproportionate number of genes associated with photosynthesis and amino acid metabolism in comparison to genes with characteristics of R-proteins or involved in secondary metabolism. The former processes may have given *S. sisymbriifolium* a bigger competitive advantage than the latter did.

## Introduction

*Solanum sisymbriifolium* (SSI), otherwise known as “litchi tomato”, “*morelle de Balbis*”, or “sticky nightshade”, is an undomesticated relative of potato and tomato. For more than a decade, SSI has been investigated as a trap-crop (a plant that attracts nematodes but kills them before they can reproduce) for nematodes such as *Globodera pallida* that normally parasitize potatoes and tomatoes (Timmermans, 2005; Dandurand and Knudsen, 2016). It is also a potential source of anti-protozoan (Meyre-Silva *et al*., 2013) and anti-molluscan (Bagalwa *et al*., 2010) metabolites. If the genetic basis for these protective processes could be identified, it might be possible to transfer these traits, either through cross-breeding or through modern transgenic technologies, from this weed to its domesticated relatives. However, while the genomes of potato and tomato have been studied extensively, spiny solanums, like SSI, have not (Yang *et al*., 2014). Only 54 SSI nucleotide sequences have been submitted to NCBI as of 2016. This ignorance about the biology and genetics of the spiny solanums could be masking a wealth of genetic resources that could be used to protect agriculturally important crops.

Most bioinformatic analyses of a species begin with the assembly and annotation of a complete genome. Once assembled, these data can be searched for genes encoding a particular protein or RNA sequence. For those working on a species that has not been studied extensively in the past, and which is only being studied now in order to conduct a limited number of experiments, whole genome sequencing can be more expensive and time consuming than can be justified. In these circumstances, alternative methods using sequencing technologies that are generally referred to as next-generation sequencing (NGS), have allowed researchers to by-pass whole genome sequencing in favor of generating a smaller database, one depleted of the silent regions of the genome and of genes that are not contributing to the phenotypes of interest. Most commonly, this is done using Illumina or 454 platforms that generate 10’s and 100’s of millions of short reads from cDNA copies of all of the mRNAs expressed during a given moment of time. Once obtained, these sequences can then be merged *in silico* into full length protein coding sequences. However, this *de novo* transcriptome can sometimes prove problematic. Short reads derived from highly conserved coding domains and repetitively organized genes can potentially be aligned and joined into chimeric assemblies that cannot be verified or removed because there is no independently sequenced genome available to serve as an extended template or scaffold to ensure that the merged sequences are indeed co-linear (Yang and Smith, 2013). A recent technical improvement, Pacific Biosciences’ single-molecule real-time (SMRT) “sequencing by synthesis” strategy, has become sufficiently accurate and attainably priced to be utilized by small research groups. The benefit of using SMRT sequencing is that it produces vastly longer reads than previous methodologies, although with lower coverage (Eid *et al*., 2009). The longer reads allow researchers to establish a transcriptome consisting of nearly complete open reading frames free of the kinds of errors possible when sequences must be assembled *in silico* from short reads (Ocwieja *et al*., 2012; Zhang *et al*., 2014).

The specific goal of the current project was to establish a four organ (bud, leaf, stem and root) *de novo* transcriptome of SSI. In doing so, we wanted to ensure that the final sequences were high-quality and consisted of genes that were biologically relevant and not artifacts of some *in silico* assembly process. This transcriptome will provide a reference library to be used in future RNA-seq experiments to identify genes for nematode and other pathogen resistances in SSI.

## Results

### Establishing a SMRT sequenced transcriptome

Before any sequencing was attempted, the genome size of SSI was estimated using flow cytometry (Supplemental Figure 1). This showed that the genome mass of SSI was approximately 4.73 pg per 2C, or 2,315 mega-base pairs per 1C. By comparison, Arumuganathan and Earle, 1991, using the same technology estimated that the tomato genome massed between 1.88 to 2.07 pg per 2C while tetraploid potato massed between 3.31 to 3.86 pg per 2C. Thus, these initial measurements gave SSI a genome size greater than tetraploid potato. Despite their unusual length (Paul and Banerjee, 2015), SSI has 24 chromosomes, like diploid potatoes and many other Solanaceae (data not shown). Due to the size of this genome, and our interest in generating a database of protein-coding genes, we elected to sequence the SSI transcriptome rather than its genome.

Generating an SSI transcriptome was done using Single-Molecule Real Time (SMRT) sequencing by PacBio Sciences (Eid *et al*., 2009) that does not need to assemble short reads into one contiguous sequence. Rather than producing only short reads, SMRT technology can provide reads up to 60,000 bp along a single molecule of DNA. This allows the capture of entire genes with one read rather than chunking it into many small bits that have to be assembled later. This sequencing strategy gives higher coverage than Sanger-based reactions like those performed on Applied Biosystems™ gene analysis instruments, and longer reads than Illumina or Roche 454.

The Iso-Seq pipeline classifies the sequences as either full-length non-chimeric (FLNC), or non-full length reads. Full length reads are those containing both 5’ and 3’ adapters, in addition to the poly(A) tail. The reads containing these parts in the expected order, i.e. 5’ adapter–poly(A)–3’ adapter, with no additional copies of these parts, are classified as non-chimeric. The FLNC reads in the present dataset were corrected with the non-full length reads using Iterative Cluster for Error correction and the Pacific Biosciences Quiver algorithm (https://github.com/PacificBiosciences/SMRT-Analysis/wiki/ConsensusTools-v2.3.0-Documentation).

In an attempt to improve our ability to detect differences in the suites of genes expressed in different parts of the plant, we generated cDNA from 4 organs; leaves, stems, roots, and unopened flower buds. We then independently carried out SMRT sequencing of all 4 samples. Finally, all corrected FLNC reads were merged *in silico* and redundancy was removed using CD-HIT-EST (Li and Godzik, 2006).

This SMRT sequencing strategy created 231,712 total corrected FLNC sequences (Table 1) using the aforementioned pipeline. These sequences had a GC content of 41.2%. CD-HIT-EST was then used to reduce the redundant sequences to sets with 100% identity. This lowered the number of sequences to 139,611 with a GC content of 41.0%. The GC content continued to decrease as the identity was reduced using CD-HIT-EST. At 80% identity, there were 32,315 sequences, with an estimated GC content of 39.7%. The decrease in GC content could be due to the methodology of CD-HIT-EST that retains the longest sequence during the reduction process. Because of this, reads that spanned untranslated regions of a transcript were expected to be favored over those only consisting of coding regions.

**Table 1:** Summary of the SSI transcriptome derived using SMRT technology. Organ sub-transcriptomes were sequenced and combined from 33,170 root, 99,924 bud, 50,825 leaf, and 47,793 stem reads. See Supplemental Figure 3 and Supplemental Figure 4 for gene ontology bins of this transcriptome.

| SMRT Assembly |  | SSI |
| --- | --- | --- |
| Total raw reads |  | 231,712 |
| Read lengths |  | 300-7883 |
| Total raw reads size (bp) |  | 362,086,346 |
| GC content |  | 41.17 |
| Transcripts | Number | 139,611 |
|  | Total length | 237,865,670 |
|  | N50 | 2,050 |
|  | Max length | 7,883 |
|  | GC content | 40.97 |
| Unigenes | Number | 41,189 |
|  | Total length | 74,642,518 |
|  | N50 | 2,158 |
|  | Max length | 7,883 |
|  | GC content | 39.90 |

In the end, we chose to work with a final SMRT dataset that had been reduced to 90% identity and consisted of a set of 41,189 sequences with a GC content of 39.9%. We judged that this estimate of the number of transcripts present in the 4 organs would most likely err on the high side, yet still retain most splice variants within a gene, as well as many paralogs and single nucleotide polymorphisms between alleles of this obligate outbreeder.

### Evidence based Quality Control of the SMRT Transcriptome

We performed an internal quality check by sequencing 45 randomly chosen clones from a cDNA library using Sanger Dye Deoxy technology (ABI 3730, Applied Biosystems). Bowtie2 (Langmead and Salzberg, 2012) was then used to find the most likely equivalent of each clone in our SSI transcriptome. A manual comparison of the Sanger-sequenced clone and the assembled transcript was done using DNA Strider (Marck, 1988). Firstly, all 45 cDNAs were found in the SSI transcriptome (Figure 1A). Secondly, only two of these SMRT-derived sequences appeared to be chimeric (Figure 1B), and based on the length of non-homologous stretches, could have been transcribed from different members of the same gene family rather than been created by misassembly. During our analysis of the SSI transcriptome, we did find entries that consisted of inverted repeats of entire gene sequences. These inverted repeats likely occurred during the preparation of the cDNA library prior to sequencing rather than during sequencing or subsequent computational processing as can be found in Illumina or 454 assemblies (Loman *et al*., 2012; Luo *et al*., 2012).

**Figure 1:**
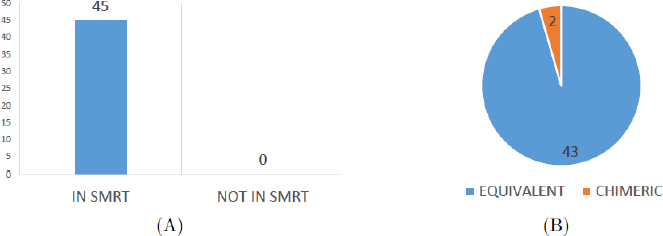
Correspondence between 45 randomly selected Sanger-sequenced SSI cDNAs and the SMRT transcriptome. A) Bowtie2 was used to determine the presence of 45 cDNA clones in the SMRT SSI transcriptome. B) Alignment of matched sequences to SMRT SSI transcriptome was performed using DNA Strider and manually evaluated as either equivalent or chimeric. All Sanger sequenced clones were found in the SMRT dataset and a small percentage were found to be chimeric. See Supplemental Figure 2 for corresponding analysis of Illumina sequenced, Velvet/Oases transcriptome.

When the SSI transcriptome was analyzed using Mercator (Lohse *et al*., 2014), it contained 38.6% unannotated sequences. This was markedly fewer than the percent unannotated sequences of either potato (50.8%) or tomato (46.4%) transcriptomes processed in the same way via Mercator (Supplemental Figure 4). Other than that, the binned profile of SSI was very similar to the published transcriptomes (Supplemental Figure 3) of these plants. This led us to believe that our transcriptome was at least of comparable quality with the working transcriptomes of these two better studied species.

PfamScan (Finn *et al*., 2008) was also used to annotate the domains of the transcriptome. This program uses HMMer (Eddy, 1998) domain annotations, and used in combination with protocols established by Sarris *et al*., 2016, allowed for the annotation of domains found in the amino acid sequences translated from the assembled transcriptome. This annotated 84.7% of the transcriptome with at least one recognizable domain. There were fewer unannotated sequences in the SSI transcriptome than in the STU and SLY transcriptomes (Supplemental Table 1). The reduced number of unannotated sequences found in the SMRT transcriptome might reflect the fact that this set had undergone a conservative reduction to 90% identity. Alternatively, the reduced number of unannotated SSI sequences could have resulted from the fact that we had only sampled the four most frequently studied organs of a “normally” growing plant, that is, plants manifesting a physiological state which has been extensively studied in numerous species, while the STU and SLY transcriptomes were compiled from plants sampled over a much broader range of life-history stages and growth conditions ranging from fruit and tuber development to exposure to biotic and abiotic stresses where the functions of many genes are still under investigation.

To further test the quality and completeness of our transcriptome, BUSCO benchmarking (Simão *et al*., 2015) was performed. The BUSCO database was established to allow researchers to assess the completeness of new genomes or transcriptomes based on the detection of a set of universal, single-copy orthologs. We found 93% intact BUSCO archetypes (889 genes) in the SSI SMRT transcriptome, 30.2% (289) of these were found in multiple copies, while an additional 2.2% (21) of the BUSCO archetypes were present in fragments, and 4.8% (46) were missing entirely (Table 2). These numbers representing genes expressed during a single growth condition of SSI, were only 4.7 percentages different from the numbers of BUSCO archetypes found in the entire SMRT sequenced genome of *A. thaliana*.

**Table 2:**
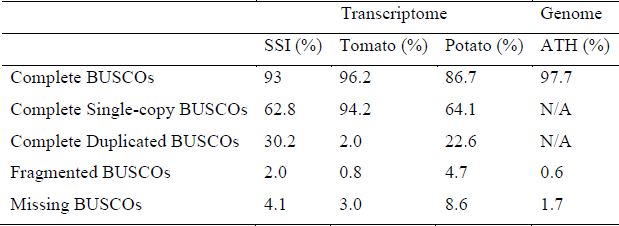
BUSCO assessment for completeness of 3 transcriptomes and one genome. The SSI transcriptome appears to be nearly complete, but contains a disproportion number of duplicated sequences. See Supplemental Table 4 for sources of datasets.

|  | Transcriptome |  |  | Genome |
| --- | --- | --- | --- | --- |
|  | SSI (%) | Tomato (%) | Potato (%) | ATH (%) |
| Complete BUSCOs | 93 | 96.2 | 86.7 | 97.7 |
| Complete Single-copy BUSCOs | 62.8 | 94.2 | 64.1 | N/A |
| Complete Duplicated BUSCOs | 30.2 | 2.0 | 22.6 | N/A |
| Fragmented BUSCOs | 2.0 | 0.8 | 4.7 | 0.6 |
| Missing BUSCOs | 4.1 | 3.0 | 8.6 | 1.7 |

SSI has the same number of chromosomes as most other Solanaceae, and does not appear to be polyploid (data not shown), yet the BUSCO analysis showed that SSI had more duplicate copy archetypes than diploid and tetraploid potato. This high number of similar sequences could point to the fact that our transcriptome has not been reduced far enough, or could be one line of evidence that SSI has undergone extensive genome duplication or hybridization in the past. This latter hypothesis was evaluated by divergent gene analysis as has been done with plants such as wheat (Krasileva *et al*., 2013). When the program Freebayes (Garrison and Marth, 2012) was run using a defined diploid setting, it output information stating there were genes that had more than 2 alleles or paralogues. We redid the analysis using defined triploid and tetraploid settings and found that even after merging sequences with more than 90% identity using CD-HIT-EST, the SSI transcriptome contained 1,348 genes with 3 distinguishable alleles or paralogues and furthermore, 44 genes with 4 distinguishable copies (Table 3). It was noteworthy that no gene had more than 4 alleles or paralogues. A simple explanation for these multiple gene variants, that would be consistent with the BUSCO analysis, was that SSI underwent a genome duplication followed by diploidization in the past and that over time, some of the duplicated loci acquired additional mutations while other loci were lost. To determine if this proposed duplication was restricted to one chromosome, or one chromosomal arm, the 44 genes with 4 alleles were mapped onto SLY chromosomes (Supplemental Table 2). There were “4–allele” genes found on 11 of 12 SLY chromosomes which indicated, assuming that genes dispersed in tomato were not linked when the two species diverged, that SSI has undergone a full genome duplication rather than a segmental duplication within one chromosome.

**Table 3:**
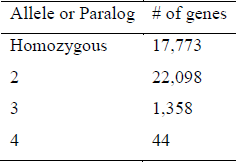
Divergent gene assessment of allele and/or paralog number in the SSI transcriptome. 4-allele genes were mapped to tomato chromosomes, see Supplemental Table 2.

| Allele or Paralog | # of genes |
| --- | --- |
| Homozygous | 17,773 |
| 2 | 22,098 |
| 3 | 1,358 |
| 4 | 44 |

Since SSI is not as well-known as other Solaneacae, we employed OrthoMCL v2.0.9 (Li *et al*., 2003) to illustrate some of the common features its gene complement showed with those of other plants. Protein sequences from our SSI transcriptome (translated using the program ESTScan (Iseli *et al*., 1999)), and protein sequences from tomato (SLY), potato (STU), eggplant (SME), *Arabidopsis thaliana* (ATH), papaya (CPA), grapes (VVI), peaches (PPE), black cottonwood (PTR), oranges (CSI), alfalfa (MTR), maize (ZMA) and rice (OSA) were merged into 45,234 orthologous groups (gene families). In this set, 6097 orthologous groups were shared by all 13 species (Figure 2), an overlap well within the range of previous studies (Yang *et al*., 2014). Each species had many additional groups that were not shown in this diagram because they were not shared with all members of this set of plants. Interestingly, even closely related species like SSI, STU, SLY, and SME had hundreds of groups not found in each other. When the annotations of the SSI unique set were compared to the full transcriptome, several functional groups showed a disproportionate increase. It is possible that these disproportionately expanded sets, that included photosynthetic genes, and genes for amino acid and vitamin metabolism (Supplemental Figure 5), diverged so much more than groups such as those for cell wall composition, and hormone and secondary metabolism, because expansion of the former traits gave SSI a competitive edge over other species in their habitat. Overall, though, there were fewer groups of genes unique to SSI than unique to STU and SME. As noted previously, this could merely reflect the fact that our data came from a single-point snapshot of only 4 organs and so would have lacked those transcripts specifically expressed during fruit and seed set, germination, senescence, abiotic stress, pathogen attacks, and numerous other stages of a plant’s lifecycle.

**Figure 2:**
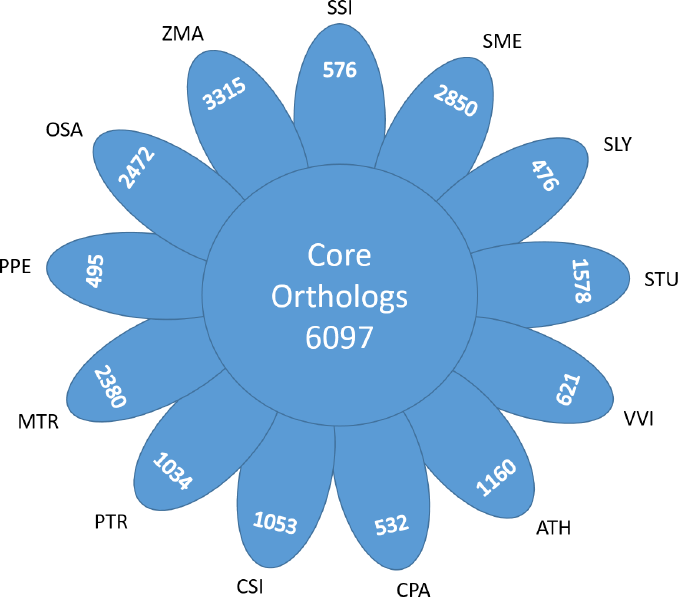
Shared and restricted orthologous genes among 13 species. All species shown here shared 6067 core orthologs. Each petal shows the number of gene groups unique to each species. Not shown are groups shared by only 2–12 species. *Solanum sisymbriifolium*, SSI; *Solanum tuberosum*, STU; *Solanum lycopersicum*, SLY; *Solanum melongena*, SME; *Arabidopsis thaliana*, ATH; *Carica papaya*, CPA; *Vitis vinifera*, VVI; *Prunus persica*, PPE; *Populus trichocarpa*, PTR; *Citrus sinensis*, CSI; *Medicago truncatula*, MTR; *Zea mays*, ZMA; and *Oryza sativa*, OSA. See Supplemental Figure 5 for gene ontology bins for the SSI unique groups, and Supplemental Table 4 for sources of datasets.

Using highly conserved orthologous genes, i.e. subunits of Rubisco, provisional phylogenies were created for nuclear-encoded and chloroplast-encoded genes using the aforementioned species (data not shown). In doing so, we concluded that nuclear SSI was most closely related to eggplant, which has been noted previously (Särkinen *et al*., 2015), while chloroplast SSI was more closely related to tomato. This dichotomy has also been seen by others (Miz *et al*., 2008) and interpreted to indicate that SSI had undergone an ancient hybridization and afterwards retained the chloroplast genome from one parent, and much of the nuclear genome from another. However, many more SSI genes will have to be compared with the genes of many more South American plants to confirm that this hybridization occurred.

### Building a snapshot of organ-associated gene expression

Since we had maintained separate cDNA pools from individual organs, it was possible to backtrack each sequence within the final transcriptome to obtain a provisional profile of gene expression throughout the plant (Figure 3A). This analysis showed that there were 8019 sequences expressed solely in buds, 4957 solely in roots, 5349 solely in leaves, 4198 solely in stems, and 7212 sequences expressed in all tissues. That left 11,538 sequences that were expressed in more than one organ but not in all 4.

**Figure 3:**
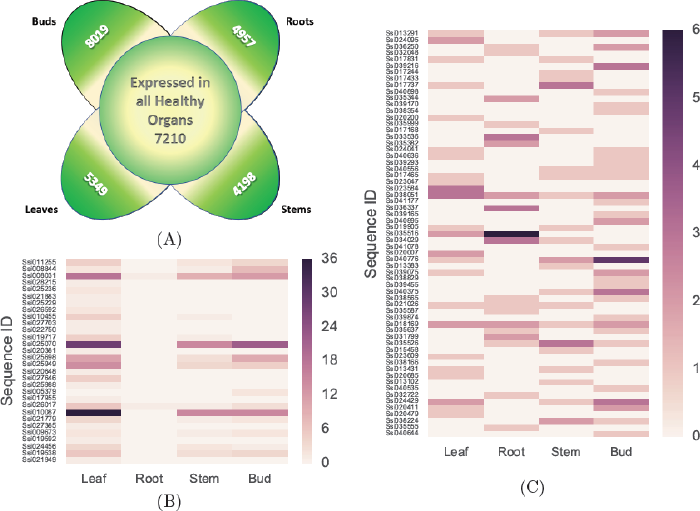
Final SMRT transcriptome sequences were backtracked through the de-redundification process to the organ sub-transcriptomes. A) Flower plot of genes expressed in all, or in only one, organ. Each petal shows the number of genes only expressed in one organ. In the center are the number of genes expressed in all 4 organs. Not shown are genes expressed in 2 or 3 organs. B) Green-tissue specificity of sequences annotated as genes involved in the light harvesting complex-I pathway via Mercator. C) Sequences annotated as putative resistance genes because they contained nucleotide binding (NB-ARC) and leucine rich repeat (LRR) domains showed varied expression patterns. As shown on the scales on the right of (B) and (C), the darker the color, the more times the sequence was found in that organ.

This backtracking allowed us to construct an expression snapshot that showed how different genes were being expressed at the time the organs were harvested. Using several in-house Python scripts, we recorded the number of reads for genes that had common annotations for several different physiological processes.

A set of light-harvesting complex genes (LHC-I) were predictably found in aerial organs with few exceptions (Figure 3B), demonstrating that the backtracking program could extract biologically useful information about sequences with specified characteristics from the merged transcriptome. In order to determine if this kind of analysis of SMRT sequences could categorize the expression of very different sets of genes, we constructed an inter-organ expression profile of genes that encoded both a leucine-rich repeat (LRR) domain together and a nucleotide binding (NB-ARC) domain (Figure 3C), a pairing frequently found in pathogen resistance genes (R-genes). This profile of R-gene prevalence in SSI, potato, and tomato indicated that there was a reduction of these genes in the SMRT transcriptome compared to the other two species (Table 4). Three of these potential R-genes were then assayed by semi-quantitative PCR (primers found in Supplemental Table 3) and quantified using a sample of cDNA from the same pool that had been sequenced, and a sample from an independently-prepared, unsequenced cDNA pool (Figure 4). In order to assess whether a SMRT data set could be a reliable indicator of gene expression, both the *in silico* and PCR measurements of gene expression were normalized in kind to an actin sequence (Ssi032526). The physical measurements of expression of two of the three genes matched the expression snapshot extremely well, but the third gene (Ssi038051) was more abundant in stems and buds than expected based on its SMRT expression snapshot. This confirms that whole transcriptome snapshots can provide a provisional picture of organ differences in gene expression, but further shows that the expression of each gene of interest needs to be verified biologically, most usefully by multiple independent tests.

**Figure 4:**
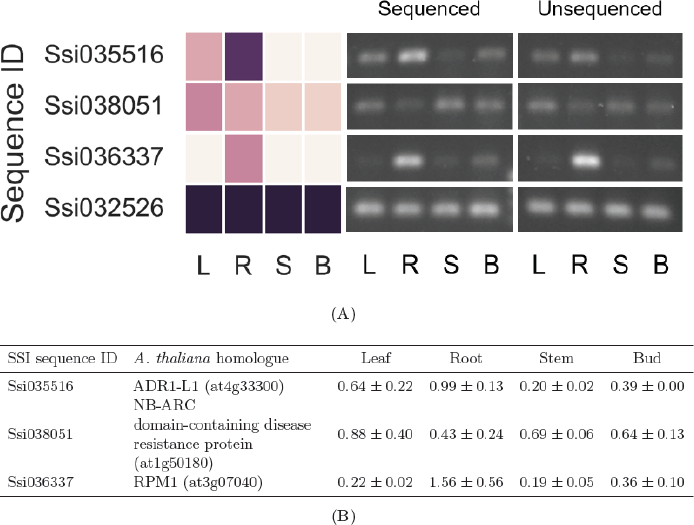
Comparison of expression of 3 putative R-gene sequences in the SMRT database to semi-quantitative PCR from 2 cDNA preparations. A) The expression of three genes with LRR and NB-ARC domains characteristic of the R-genes and an actin isoform is shown in the heat map at the left and compared on the right to semi-quantitative PCR of those same genes in two independently prepared cDNA pools, one from the pool used to generate the transcriptome (Sequenced) and one prepared independently, and not used to make the transcriptome (Unsequenced). B) The expression of each PCR product from each pool was quantified and then normalized to the expression of an actin isoform (Ssi032526). Data (biological replicates, n=2) are represented as mean ± STD. See Supplemental Table 3 for primers used.

**Table 4:**
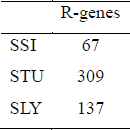
R-gene profile of potato (STU), tomato (SLY), and the SSI transcriptome. The SSI database had fewer assigned R genes (based on the presence of nucleotide-binding domains and leucine-rich repeats within the same open reading frame) than either SLY or STU genomes. Refer to Supplemental Table 1 for full domain annotation statistics.

## Discussion

The creation of a *de novo* transcriptome necessitates massive amounts of follow-up analyses, both *in silico* and biologically, to estimate its reliability.

We initially employed both Illumina and 454 sequencing (data not shown) in order to compensate for the different kinds of errors to which each method was prone (Luo *et al*., 2012). Screening this assembly with genes randomly selected from an SSI cDNA library revealed that 20% of these genes failed to match any of the assembled sequences in this database (Supplemental Figure 2A), and of those that matched, 40% appeared to be chimeric (Supplemental Figure 2B). In contrast, all of these cDNAs were found in our SMRT sequenced transcriptome and few were patently chimeric (Figure 1).

A number of factors are known to exasperate misassembly including the presence of large gene families and of repeatedly occurring kmers in the dataset (Moreton *et al*., 2015). Even though we did not sequence the SSI genome, we found 4 lines of evidence indicating that it might be complex enough to pre-dispose our transcriptome to these kinds of assembly mistakes. First, the nuclear DNA content of SSI was larger than most diploid Solanaceae, roughly the same size as a tetraploid potato (Supplemental Figure 1). Second, divergent gene analysis indicated that the SSI transcriptome was unusually complex and contained 3 and 4 distinguishable alleles for many genes (Table 3). Third, there were only 67 putative R-genes, that is, genes containing a nucleotide binding domain (NB-ARC) and a leucine-rich repeat domain (LRR), in the SMRT sequenced dataset compared to the 309 in STU, and 137 in STU (Table 4). Finally, an unusually high percentage of the BUSCO gene set were present in multiple copies in SSI even though our transcriptome could only consist of a portion of all the genes that are likely to be encoded in its DNA (Table 2). One model consistent with these 4 facts was that SSI had, sometime in the past, undergone a partial or complete genome duplication. Over time, as diploidy was re-established, some of the duplicated alleles or paralogues diverged, while others were lost. Nevertheless, enough of the expanded gene families remained to confound the alignment programs that tried to differentiate between their members. While these kinds of errors might be correctable with the use of other assembly programs, we chose, instead to create an assembly-independent transcriptome using SMRT technology.

At the moment, SMRT technology does not provide the sequence coverage or depth that can be obtained with Illumina or 454 sequencing. In order to increase our chances of sampling uncommon organ-specific transcripts, we prepared independent cDNA pools from 4 organs of the plant. Using an in-house script (https://github.com/AlexWixom/Transcriptome_scripts), we were able to increase the value of the final library by generating expression snapshots for genes of interest in each organ. These expression snapshots are no substitute for a more thorough RNA-seq study, but they do provide a preliminary assessment of a plant’s biology at the time of harvest. Using these snapshots, we recognized different patterns of expression of individual LHC-1 genes (Figure 3B) within the photosynthetic parts of the plant. We also saw that 2 of the 3 R-genes re-examined by PCR showed the same expression pattern in two independent RNA and cDNA preparations as found in the transcriptome itself (Figure 4). Thus, in the absence of RNA-seq studies or experimental evidence for the role of a specific locus, this kind of library assembly could be used to direct researchers to the subset of R-genes most likely responsible for the resistance in a given organ.

R-genes coding for recognition proteins are commonly perceived as sentinels that are awaiting activation by molecules introduced during infection (Jones and Dangl, 2006). Therefore, the reduced number of R-genes found in SSI (67), compared to both potato (309), or tomato (137) was unexpected. This discrepancy could be explained in any one of several ways. First, SSI might be using proteins with novel domain structures in place of classic R-genes. Second, sequencing depth might simply have been inadequate to capture all R-genes that were actually being expressed at low levels. Finally, SSI could be relying on rapidly inducing transcription of R-genes after an infection has occurred. Any one of these hypotheses is worthy of continuing analysis.

With this transcriptome as an example, we have established a protocol that opens the door to further genetic mining of previously uncharacterized species. The completeness of our database indicates that *de novo* transcriptomes not only provide an economical and time-saving way to study a new species, but can also provide expression data that could not be gleaned from a genomic sequence.

## Materials and Methods

### Plant and Culture Conditions

*S. sisymbriifolium* (SSI) seeds obtained from C. Brown (USDA-ARS, Prosser WA) were germinated in soil. Nodes from a single plant were sterilized for 20 min using 10% NaClO with 0.05% Tween20. Plant material was then washed 3x with sterile distilled H2O and put into 120 mL baby food jars containing standard Murashige and Skoog salts, pH 5.6, 3% sucrose, 0.7% agar, 100 μg mL^−1^ myo-inositol, 2.0 μg mL^−1^ glycine, 1 μg mL^−1^ thiamine, 0.5 μg mL^−1^ pyridoxine, and 0.5 μg mL^−1^ nicotinic acid. A single plant was chosen as the progenitor of all of the plants used in this study. All of its descendants were maintained at 25°C in 16 h light, and subcultured vegetatively every 4 wk. Over the course of the project, rooted clones with at least 4-6 leaves were put into 2 L of hydroponic medium (Yoshida *et al*., 1976), referred to here as Fake Field. Each container was diffusely aerated through an aquarium stone, maintained at constant volume by the addition of distilled water, and emptied and refilled with fresh hydroponic medium every 7 d. Hydroponic containers were maintained at 22°C, 16 h light with an irradiance level of 0.0006 W m^−2^. Illumination was provided by GE Lighting Fluorescent lamps (13781, F96T12/CW/1500). After a 2 wk lag-time, plants began producing 1-3 new leaves each wk, and flowered continuously afterwards. All experiments were performed on plants that had not been infected or wounded in any way previously.

### RNA extraction

RNA was extracted from SSI bud, stem, leaf, and root and infected root organs adapted from the protocol in Casavant *et al*., 2017. Adaptions included use of a coffee grinder to homogenize tissue with the addition of dry ice to maintain RNA integrity.

### Genome size estimation by Flow Cytometry

Healthy green leaf tissues were collected from SSI plantlets growing in vitro. Roughly 1 cm^2^ (0.01g or less) of leaf was chopped in 1mL ice cold LB01 buffer for 1.5-2 min (Doležel *et al*., 1989). The LB01 buffer contained 50μg mL^−1^ RNase stock and 50μg mL^−1^ propidium iodide (25% PI stock in DMSO) per mL of LB01. Each sample was chopped with a fresh razor blade in a clean Pyrex petri plate. The finely chopped suspension was then filtered through a 50μm nylon mesh filter (Partec 04-0042-2317). This filtered suspension was kept in the dark at 4°C for between 15-90 min before it was analyzed.

Genome size estimations were made using a BD FACSARIA Flow Cytometer (IBEST Imaging Core, University of Idaho, Moscow, ID, USA). A green laser at 488nm was used to excite the propidium iodide stained cells and was then collected in the PE-A channel. Thresholds for PE-A were set at 1,000 and FSC at 500. The voltages were set so the major peak (2C) of the SSI samples were near 50,000 on the linear scale. Four suspensions were made from separate donor plants once a day for three consecutive days. Two replicates of two external standards were also used daily in addition to the 4 SSI samples. External standards included *Solanum lycopersicum cv.* Stupicke polni tyckove rane (2C= 1.96 pg DNA) and *Glycine max* cv. Polanka (2C= 2.50 pg DNA) (Doležel *et al*., 1992; Doležel *et al*., 1994) which were chosen because their genome sizes were in the expected range of SSI. One repetition of internal standards was run using tomato and soybean. DNA content was estimated using the equation described by Doležel *et al*., 2007.

### Library Preparation for Iso-Seq

SMRT library preparation and sequencing were performed by the National Center for Genome Resources (Santa Fe, New Mexico). The Iso-Seq libraries for four organs, root, stem, leaf and bud, were prepared for Isoform Sequencing (Iso-Seq) using the Clontech SMARTer PCR cDNA Synthesis Kit and the BluePippin Size Selection System protocol as described by Pacific Biosciences (https://goo.gl/ij71Hh) with the following modifications. For cDNA conversion, 3 μg of total RNA was put into each Clonetech SMARTer reaction. From the PCR optimization procedure specified in the protocol, it was determined that 14 cycles of PCR would be sufficient for amplification of each organ’s cDNA. Amplification was followed by size selection on each sample to obtain three size bins (0.5-2 kb, 1.5-3 kb and 2.5-6 kb) using the Blue Pippin (Sage Science, Beverly, Massachusetts) instrument. The amplified and size selected cDNA products were made into SMRTbell Template libraries per the Isoform Sequencing protocol referenced above. Libraries were prepared for sequencing by annealing a sequencing primer (component of the SMRTbell Template Prep Kit 1.0) and then binding polymerase to this primer-annealed template. The polymerase-bound template was bound to MagBeads (P/N 100-125-900) (https://goo.gl/wdZErU) and sequencing was performed on a PacBio RS II instrument. 12 v3 SMRTcells were run for the root tissues, 14 for the leaf tissues, 9 for the stem tissues, and 12 for the bud tissues for a total of 47 SMRTcells (Pacific Biosciences, P/N 100-171-800). The libraries from each organ were separately sequenced using P6C4 polymerase and chemistry and 240-minute movie times (Pacific Biosciences, P/N 100-372-700, P/N 100-356-200).

### Single Molecule Real Time Sequencing

All SMRT cells for a given organ were run through the Iso-Seq pipeline included in the SMRT Analysis software package. First, reads of insert (ROIs, previously known as circular consensus sequences or CCS) were generated using the minimum filtering requirement of 0 or greater passes of the insert and a minimum read quality of 75. This allowed for the high yields going into subsequent steps, while providing high accuracy consensus sequences where possible. The pipeline then classified the ROI in terms of full-length, nonchimeric and non-full length reads. This was done by identifying the 5’ and 3’ adapters used in the library preparation as well as the poly(A) tail. Only reads that contained all three in the expected arrangement and did not contain any additional copies of the adapter sequence within the DNA fragment were classified as full-length non-chimeric copies. Finally, all full-length non-chimeric reads were run through the Iterative Clustering for Error correction algorithm then further corrected by the Pacific Biosciences Quiver algorithm (https://github.com/PacificBiosciences/cDNA\_primer/wiki/Understanding-PacBio-transcriptome-data). Once the Iso-Seq pipeline result was available for each organ, the results were combined into a single data set and redundant sequences were removed using CD-HIT-EST (Li and Godzik, 2006).

### Illumina Sequencing for divergent gene analysis

Extracted total RNA from each previously stated organ was sent to Eurofins Genomics (Ebersberg, Germany) for library preparation and sequencing. Prior to library preparation, quality control (QC) was performed on individual tubes of RNA and equal aliquots of each preparation were blended into one pool. The Illumina library cDNA was prepared using randomly-primed first and second strand synthesis, followed by gel sizing and PCR amplification. The library was then physically normalized and found to have insert sizes of 250-450 bp.

Illumina sequencing was performed on a MiSeq v3 2×300. The read sequences were clipped using Trimmomatic, version 0.32 (Bolger *et al*., 2014), and bases with a Phred score < 20 were removed. Trimmed reads shorter than 150 bp were removed; this step could remove none, one, or both mates of a read-pair. Digital normalization was applied to the Illumina reads in order to reduce redundant information present in these large datasets. A coverage cutoff of 30 and a kmer size of 20 decreased the data to 28% of the initial 31,310,146 reads. BWA (Li and Durbin, 2009) was used to map read pairs to the *Solanum tuberosum cultivar* Desiree chloroplast genome (GenBank accession DQ38616.2). Only unmapped reads were retained.

### Annotation of Sequences

Mercator sequence annotation was performed using the TAIR, PPAP, KOG, CDD, IPR, BLAST CUTOFF of 80, and ANNOTATE options (Lohse *et al*., 2014).

### Annotating Protein Domains of Translated Sequences

PfamScan (Finn *et al*., 2009) was run on the SSI transcriptomes following protocols set forth by Sarris *et al*., 2016.

### Biological Quality Check of *in silico* Sequences

45 clones from a cDNA library (Express Genomics, Average insert size=1 kb, Vector= pExpress 1) were randomly selected and sequenced via Sanger Dye-Deoxy DNA Sequencing (ABI 3730). These sequences were then aligned to the transcriptomes using Bowtie2 (Langmead and Salzberg, 2012) set for local alignment and best hit only. These aligned sequences were then manually compared for possible chimeric features.

### Evolutionary Comparison of SSI to 13 Other Species

The evolutionary clustering and comparison protocols were adapted from those set out in Yang *et al*., 2014. See Supplemental Table 4 for species used and online download sources.

### Divergent Gene Analysis to determine Ploidy

Phasing of the SMRT transcriptome was completed using unassembled Illumina sequences adapted from protocols established by Krasileva *et al*., 2013, with the addition of an in-house Python script to quantify single-nucleotide polymorphisms present per sequence (https://github.com/AlexWixom/Transcriptome_scripts/freePloidy.py).

### Creation of Expression Snapshots Using only SMRT Sequences

In-house Python scripts were used to backtrack final transcriptome sequences to each organ using CD-HIT-EST cluster files (https://github.com/AlexWixom/Transcriptome_scripts).

### Expression Snapshot Validation

Sequence specific oligonucleotides were designed for several genes that were then used to obtain semi-quantitative PCR expression snapshots on the same cDNA used to obtain our SSI transcriptome (referred to as the “Sequenced” sample), as well as on a second cDNA pool prepared from RNA collected from independently grown plants (referred to as the “Unsequenced” sample). PCR fragment bands were quantified with a local background subtracted and normalized to actin (following the procedure established by Casavant *et al*., 2017). The primers for these genes can be found in Supplemental Table 3 with the proposed gene description.

### Accession Numbers

The SMRT sequenced transcriptome has been deposited at DDBJ/EMBL/GenBank under the accession GGFC00000000. The version described in this paper is the first version, GGFC01000000.

## Data availability

File S1 contains detailed descriptions of all supplemental files. Sequence data are available at GenBank and the accession numbers are GGFC00000000. In-house code used to generate data can be found at https://github.com/AlexWixom/Transcriptome_scripts.

## Author Contributions

NCC prepared many of the RNA samples, and all of the PCR analyses. AQW performed all remaining research, statistical analysis, and figure assemblies. AQW and ABC contributed to the experimental design. AQW wrote the manuscript with assistance from ABC. JCK, FX, and L-MD provided equipment and facilities, bacterial clones and plant material, and assisted in data interpretation. All authors have read and approved the manuscript.

## Acknowledgements

The authors would like to thank Dr. R. Tripepi for providing us with vegetatively propagated *S. sisymbriifolium*. We are also extremely grateful to the National Center for Genomics Research for preparation and sequencing of SMRT libraries. This research was funded by the Idaho Potato Commission, the Northwest Potato Research Consortium, and USDA-APHIS Farm Bill 10201, 10007.

